# Association analysis of loci implied in “buffering” epistasis

**DOI:** 10.1101/637579

**Authors:** Andrés Legarra, Zulma G. Vitezica, Marina Naval-Sánchez, John Henshall, Fernanda Raidan, Yutao Li, Karin Meyer, Nicholas J. Hudson, Laercio R. Porto-Neto, Antonio Reverter

**Affiliations:** INRA/INPT, UMR 1388 GenPhySE, F-31326 Castanet-Tolosan, France; CSIRO Agriculture & Food, 306 Carmody Rd., St. Lucia, Brisbane, QLD 4067, Australia; Cobb-Vantress Inc., Siloam Springs, Arkansas 72761-1030, USA; Animal Genetics and Breeding Unit, University of New England, Armidale, NSW 2351, Australia; School of Agriculture and Food Sciences, The University of Queensland, Gatton, QLD 4343, Australia

**Keywords:** Epistasis, genome-wide association studies, beef cattle

## Abstract

The existence of buffering mechanisms is an emerging property of biological networks, and this results in the possible existence of “buffering” loci, that would allow buildup of robustness through evolution. So far, there are no explicit methods to find loci implied in buffering mechanisms. However, buffering can be seen as interaction with genetic background. Here we develop this idea into a tractable model for quantitative genetics, in which the buffering effect of one locus with many other loci is condensed into a single (statistical) effect, multiplicative on the total (statistical) additive genetic effect. This allows easier interpretation of the results, and it also simplifies the problem of detecting epistasis from quadratic to linear in the number of loci. Armed with this formulation, we construct a linear model for genome-wide association studies that estimates, and declares significance, of multiplicative epistatic effects at single loci. The model has the form of a variance components, norm reaction model and likelihood ratio tests are used for significance. This model is a generalization and explanation of previous ones. We then test our model using bovine data: Brahman and Tropical Composite animals, phenotyped for body weight at yearling and genotyped up to ∼770,000 Single Nucleotide Polymorphisms (SNP). After association analysis and based on False Discovery Rate rules, we find a number of loci with buffering action in one, the other, or both breeds; these loci do not have significant statistical additive effect. Most of these loci have been reported in previous studies, either with an additive effect, or as footprints of selection. We identify epistatic SNPs present in or near genes encoding for proteins that are functionally enriched for peptide activity and transcription factors reported in the context of signatures of selection in multi-breed cattle population studies. These include loci known to be associated with coat color, fertility and adaptation to tropical environments. In these populations we found loci that have a non-significant statistical additive effect but a significant epistatic effect. We argue that the discovery and study of loci associated with buffering effects allows attacking the difficult problems, among others, of release of maintenance variance in artificial and natural selection, of quick adaptation to the environment, and of opposite signs of marker effects in different backgrounds. We conclude that our method and our results generate promising new perspectives for research in evolutionary and quantitative genetics based on the study of loci that buffer effect of other loci.

## INTRODUCTION

Epistasis is a biological phenomenon that is increasingly being given attention in genetics. One of the biological phenomena in which epistasis is likely implied is “buffering” (Visser *et al.* 2003; Flatt 2005), a mechanism that would allow buildup of robustness through evolution (see (Flatt 2005) for examples). A known example is chaperones (Visser *et al.* 2003; Kitano 2004). Loci implied in buffering would mitigate heritable perturbations. For instance, for a trait with intermediate optima, too high total genotypic values would not be expressed. The existence of buffering mechanisms is an emerging propriety of networks (Mackay 2014) and therefore, because biochemical and gene networks are pervasive in nature, buffering loci must exist. Moreover, the existence of segregating (not fixed) buffering epistatic loci would explain several phenomena that are not well understood: environmental robustness, release of additive variance after disturbing events, (Visser *et al.* 2003; Flatt 2005), maintenance of genetic variance in selected populations previously under stabilizing selection (Gimelfarb 1989), and opposite signs of GWAS associations in different populations (Huang *et al.* 2012).

There are so far no explicit methods to detect loci implied in buffering mechanisms. However, buffering can be understood as interaction with genetic background (Visser *et al.* 2003), and methods to detect epistasis against genetic backgrounds have been proposed (Jannink 2007). There is, in addition, increasing evidence of dependency of gene substitution effects in genetic background (Hansen 2013). More recently, a method that implicitly detects loci in interaction with genetic backgrounds has been presented (Crawford *et al.* 2017). However, in neither of these cases the connection with buffering mechanisms has been explicitly shown or put forward.

In this work, we present for the first time, to our knowledge, a formal quantitative genetics framework for the phenomenon of “buffering epistasis”, derive efficient methods for genome-wide association studies, and perform association analyses for buffering loci in two real tropical cattle populations. Next, we present the biological discoveries following these analyses, showing that significant hits in the association analysis for buffering epistasis have been reported previously either as having an additive effect or as harboring selection signatures for tropical adaptation.

## MATERIAL AND METHODS

### Biometrical model

First, we start our development with a rather general model for epistasis. Then, we show how part of the epistatic variability in this general model can be reformulated as a sum of terms of a buffering action of the “buffering” epistatic locus towards the effect of all other loci, whereas the rest of the epistatic variation is ignored. Then we reformulate the sum of buffering actions in terms of a single buffering effect and overall additive genetic value. Armed with this formulation, we describe two methods of analysis, one exact and one approximate. We generally follow existing notations (Mäki-Tanila and Hill 2014).

Consider the total genotypic effect of one individual as the sum of additive (statistical) effects of all loci (*α*_*j*_ at locus *j*), plus all possible additive by additive (statistical) interactions ((*αα*)^*ij*^ for the pairwise interaction between loci *i* and *j*):

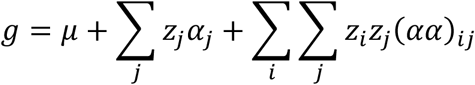

Note that these statistical pairwise interactions capture the effect of *functional* pairwise and higher-order interactions (Mäki-Tanila and Hill 2014). Gene content indicators *z*_*i*_ are centered with respect to the population, so *E*(*g*) = *μ, E*(Σ_*i*_ *z*_*i*_*α*_*i*_) = 0 and *E*(Σ_*i*_ Σ_*j*_ *z*_*i*_*z*_*j*_(*αα*)_*ij*_) = 0 across the population. The model so composed has *n* additive and *n*^2^ additive by additive effects, and they are orthogonal by construction (Cockerham 1954; Álvarez-Castro and Carlborg 2007). Consider Example in Table 1. Allele B is a “shrinker” or “bufferer” allele whereas allele b is a “magnifier” allele.

**Table 1.** Effect of buffering locus B/b on additive locus c/C.

|  | CC | Cc | cc |
| --- | --- | --- | --- |
| BB | 0 | 0.35 | 0.70 |
| Bb | 0 | 1.05 | 2.10 |
| bb | 0 | 3.15 | 6.30 |

A buffering effect can be seen as a multiplication of the additive effect of locus *j* (locus C/c in Table 1) by a quantity determined by its interaction with (buffering) locus *i* (locus B/b in Table 1), in other words, (*αα*)^*ij*^ = *k*_*ij*_*α*_*j*_. In Table 1, *k*_*ij*_ = 3. For instance, the effect of allele “c” in a genetic background “bb” is to increase 3.15 units per one unit of gene content, whereas in a “BB” background the increase is only of 0.35 units.

In our work, we assume that *k*_*ij*_ (the buffering effect of locus *i* on locus *j*) can be approximated, for all pairs of locus *i* with other loci *j*, by a locus-specific *k*_*i*_ value. In other words, for a given locus *i*, the *n* interactions (*αα*)^*ij*^ can be approximated by (*αα*)^*ij*^ ≈ *k*_*i*_*α*_*j*_. Alternatively, the value *k*_*i*_ can be seen as the regression coefficient of the equation (*αα*)^*ij*^ = *k*_*i*_*α*_*j*_ + *ϵ*_*ij*_. This simplification allows a reduction in number of parameters (from *n*^2^ interactions to *n* buffering effects), and, more important, allows focusing on individual buffering loci instead of pairs of loci. In other words, we will find loci which tend to buffer in the same manner across genome.

Thus, for the purpose of detecting buffering epistatic loci, we model the total genotypic value (ignoring remaining epistatic actions) of an individual as

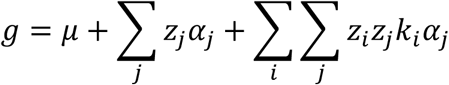

Which after some algebra and because *u* = Σ_*j*_ *z*_*j*_*α*_*j*_, becomes

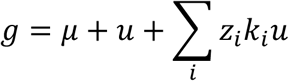

Where *u* stands for the (statistical) additive genetic value, also known as breeding value in animal genetics. For GWAS purposes, we will use a model considering only the *i*-th loci:

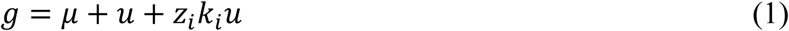

Interestingly, the term *z*_*i*_*k*_*i*_*u* can be seen as an interaction of locus *i* with all genome, as described by (Visser *et al.* 2003; Flatt 2005). This model was presented by Jannink (2007) as modelling interaction of locus with all genome, but without stating that it is in fact modelling buffering epistatic action. The same model was presented by Crawford et al. (2017)) without formalizing the kind of epistatic action, and (in particular) lacking an orthogonal model.

A feature of model in (1) is that because *E*(*u*) = 0, the observed additive effect of locus *i* changes sign in the extremes of the distribution of *u*. Imagine for instance that the buffering effect of locus *i* (B/b) is *k*_*i*_ = −0.2 and *p* = *freq*(*B*) = 0.6. For an individual with *u* = 20 and carrier of *BB* genotype, *z*_*i*_ = 2 − 2*p* = 0.8, the epistatic effect is negative: *k*_*i*_*z*_*i*_*u* = −0.2 × 0.8 × 20 = −3.2, and the total genotypic value is *g* = 20 − 3.2 = 16.8. Similarly, for an individual with *u* = 0, the epistatic locus has no effect (there is no buffering); for an individual with *u* = −20, the epistatic effect is positive and *increases* total genotypic value. In all cases, carriers of the “BB” genotype are regressed towards 0.

Plugging (1) into a linear model, a GWAS model would be

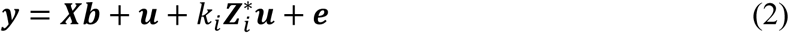

Where ***y*** are quantitative phenotypes of interest, ***b*** is the fixed effects vector (e.g. herd-sex-year contemporary group), ***X*** is a design matrix relating records to fixed effects, 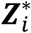 is a matrix whose diagonal contains ***z***_*i*_, the coding of the different genotypes at locus *i.* Note that the term 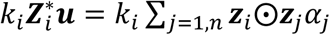 is equivalent to Eq. 1 in Crawford et al. (2017) and their terms *α*_*j*_ are equivalent to our terms *k*_*i*_*α*_*j*_. However, they do not present their model in terms of buffering, and their matrices *x* (*z* in our notation) are not centered, which leads to lack of orthogonality of their model (Álvarez-Castro and Carlborg 2007; Vitezica *et al.* 2017).

Our model in (2), is not usable because both the terms *k*_*i*_ and ***u*** implied in the regression are unknown. However, the epistatic component 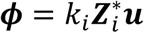 defines a covariance matrix for ***ϕ*** in the *i*-th locus:

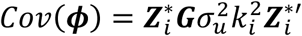

which suggests a linear model with the form ***y*** = ***Xb*** + ***u*** + ***ϕ*** + ***e***, with covariance as above (Jannink 2007; Crawford *et al.* 2017). Unfortunately, GWAS tests with this formulation imply computing and inverting *Cov*(***ϕ***) matrix at each locus (which is computing intensive) and can result in lack of convergence (Crawford *et al*., 2017). We instead propose an equivalent formulation that uses

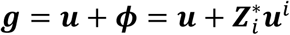

where ***u***^*i*^ = *k*_*i*_***u***. We have therefore defined a random effect, ***u***^*i*^, which multiplies real values given by the covariable 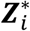. This is known as a reaction norm or random regression model (Laird and Ware 1982; Schaeffer 2004). Using this formulation, there are two additive genetic traits in this model: a general additive trait ***u*** with variance 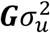, where G is a relationship matrix (Wright 1922; VanRaden 2008) and a transformation of the buffering action of locus *I* into another additive trait: 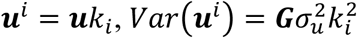. The joint covariance matrix is:

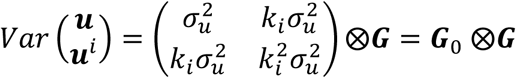

where 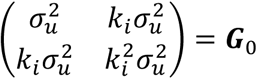 is a non-full rank matrix because ***u***^*i*^ = ***u****k*_*i*_, and ⊗ indicates the Kronecker product. Thus the final linear model, considering the epistatic interaction of locus *i* with all other loci, is

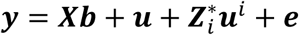

with 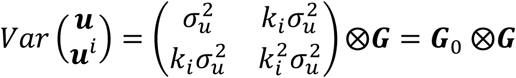. This model can be used in an exact method as described below.

### Exact maximum likelihood method

The exact method proceeds by likelihood ratio test of the two-alternative hypothesis, using random regression with the Model *H*_1_:

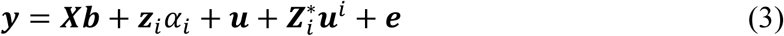

and a simpler model excluding random regression with the Model *H*_0_:

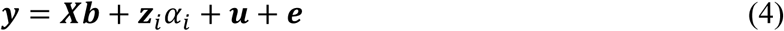

Where the actual parameter being tested is 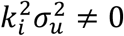. The regression on gene content ***z***_*i*_*α*_*i*_ corrects for eventual statistical additive (not epistatic) effects of locus *i*. Parameters are estimated by REML (Patterson and Thompson 1971).

After fitting the two models, the likelihood ratio test of the competing models is distributed as a mixture of 0 and 1 degrees of freedom chi-square, from which *P*-values can be obtained. In addition, from the estimated covariance matrix ***G***_0_ estimated under *H*_1_, the estimated buffering epistatic effect can be obtained as:

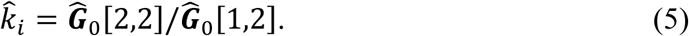

Contrary to Crawford *et al.* (2017) matrix ***G*** has to be computed and inverted only once, because inclusion of the *i*-th locus has zero influence on the result (Gianola *et al.* 2016) and the matrix 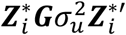 is never explicitly computed. This results in great savings of computing time. Matrix ***G***_0_ is a non-full rank matrix, which slows down convergence. An easy solution is to use a reduced rank model fitting one principal component (Meyer and Kirkpatrick 2005) as implemented in Wombat (Meyer 2007). Convergence takes a few iterations in this case compared to hundreds using a standard REML algorithm. We conceived two approximate methods that will not be reported here but whose description can be found at (Reverter *et al.* 2018).

### Animals, phenotypes and genotypes

Animals, phenotypes and genotypes used in this study were a subset of those used in Raidan *et al.* (2018). In brief, we used data of 2,111 Brahman (BB) and 2,550 Tropical Composite (TC) cows and bulls genotyped using either the BovineSNP50 (Matukumalli *et al.* 2009)) or the BovineHD (Illumina Inc., San Diego, CA) that includes more than 770,000 SNP. Animals that were genotyped with the lower density array had their genotypes imputed to higher density as described previously by Bolormaa *et al.* (2014). SNPs were mapped to the ARS-UCD1.2 bovine genome assembly. After selecting autosomal SNP with minor allele frequency (MAF) > 1%, we retained 651,253 SNPs for BB and 689,818 for TC. We used body weight at yearling (YWT) as the phenotype of interest. The average, minimum and maximum YWT (kg) were 227.7, 115 and 353 kg for BB; and 247.07, 120.5 and 394.5 kg for TC. Moreover, the average, minimum and maximum age at YWT was 360, 302 and 416 days for BB; and 361, 319 and 403 days for TC.

### False discovery rate (FDR)

Following Bolormaa et al. (2014) and with equivalent original derivations from Storey (2002), FDR was calculated as:

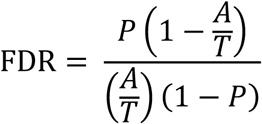

Where *P* is the *P*-value tested, *A* is the number of SNP that were significant at the *P*-value tested, and *T* is the total number of SNP tested (*T* = 651,253 and 689,818 for BB and TC, respectively).

### Implementation

We implemented the analyses using shell scripts to manipulate the data and Wombat (Meyer 2007) for the REML analyses. Markers were analyzed in parallel in the exact analyses in the Genotoul Toulouse bioinfo platform; wall clock computing time was approximately 4 days for the exact analysis run in parallel, and a few minutes for the fast approximate one (see Appendix).

### Data Availability Statement

The minimal raw data is the SNP genotype data, phenotypes and metadata (eg. sex, herd, year) for 4,661 cattle and 729,068 SNP genotypes. These raw data are part of the Beef CRC project (http://www.beefcrc.com/) and are co-owned with Meat and Livestock Australia, and can be made available upon reasonable request and subject to the agreement of the owners. Any parties seeking access to the raw data should contact Dr. Antonio Reverter ( or +61732142392).

## RESULTS

The exact method took ∼4 day in a parallel cluster running ∼100 process simultaneously. Computation time for a single marker consist of roughly 1 minute. Note that if a medium density chip had been used (50,000 markers instead of ∼600,000), computational times divide by an order of magnitude. There is considerable room for improvement of the computational methods, because most of the time is spent reading and manipulating text files.

### RESULTS – Brahman (BB) and Tropical Composite (TC) Populations

Table 2 presents the number of significant SNP and FDR at various *P*-value thresholds. At any given *P*-value, the number of significant SNP was lower in BB than in TC. As a result, the FDR was lower in TC than in BB for a given *P*-value. For instance, at *P*-value < 0.0001 the FDR was 9.83% and 4.38% for BB and TC, respectively. The higher number of epistatic SNPs identified in the TC compared to the BB population was attributed to the distinct allele frequencies observed in the two populations. Across all SNPs, the first (reference) allele was found to be either mostly absent (reference allele frequency near 0) or nearly fixated (reference allele frequency near 1) in the BB population, while intermediate allele frequencies (highly-polymorphic and hence more informative SNP) were predominant in the TC population (Figure 2 in (Reverter *et al.* 2017)). In other words, the distribution of allele frequencies in BB are U-shaped.

**Table 2.** Number of significant epistatic SNP (N) and false discovery rate (FDR) at decreasing levels of *P*-value for the Brahman and Tropical Composite populations.

| *P*-value | Brahman |  | Tropical Composite |  |
| --- | --- | --- | --- | --- |
|  | N | FDR, % | N | FDR, % |
| < 0.05 | 66,789 | 46.06 | 87,545 | 36.21 |
| < 0.01 | 20,542 | 31.01 | 30,803 | 21.61 |
| < 0.005 | 12,503 | 25.67 | 19,831 | 16.98 |
| < 0.001 | 3,679 | 17.62 | 6,972 | 9.80 |
| <0.0005 | 2,186 | 14.85 | 4,484 | 7.64 |
| <0.0001 | 662 | 9.83 | 1,571 | 4.38 |
| <0.00005 | 384 | 8.47 | 1,028 | 3.35 |
| <0.00001 | 91 | 7.15 | 342 | 2.02 |

**Table 3.**
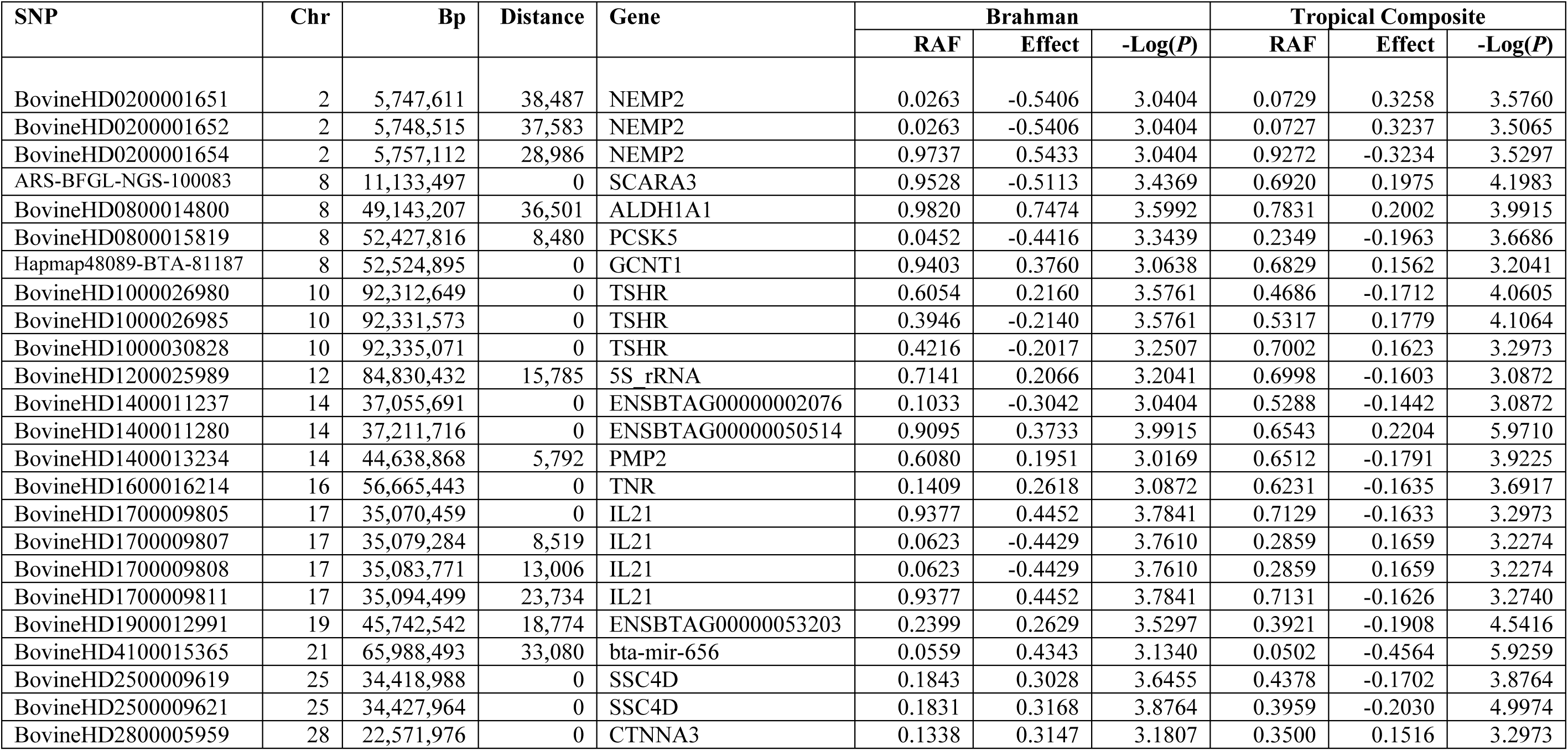
Annotation of SNP with significant (*P*<0.001) epistatic effect in both populations (Brahman and Tropical Composite) including genome position, distance to nearest gene, gene, reference allele frequency (RAF), estimated effect and significance (-Log(*P*)).

**Figure 1.**
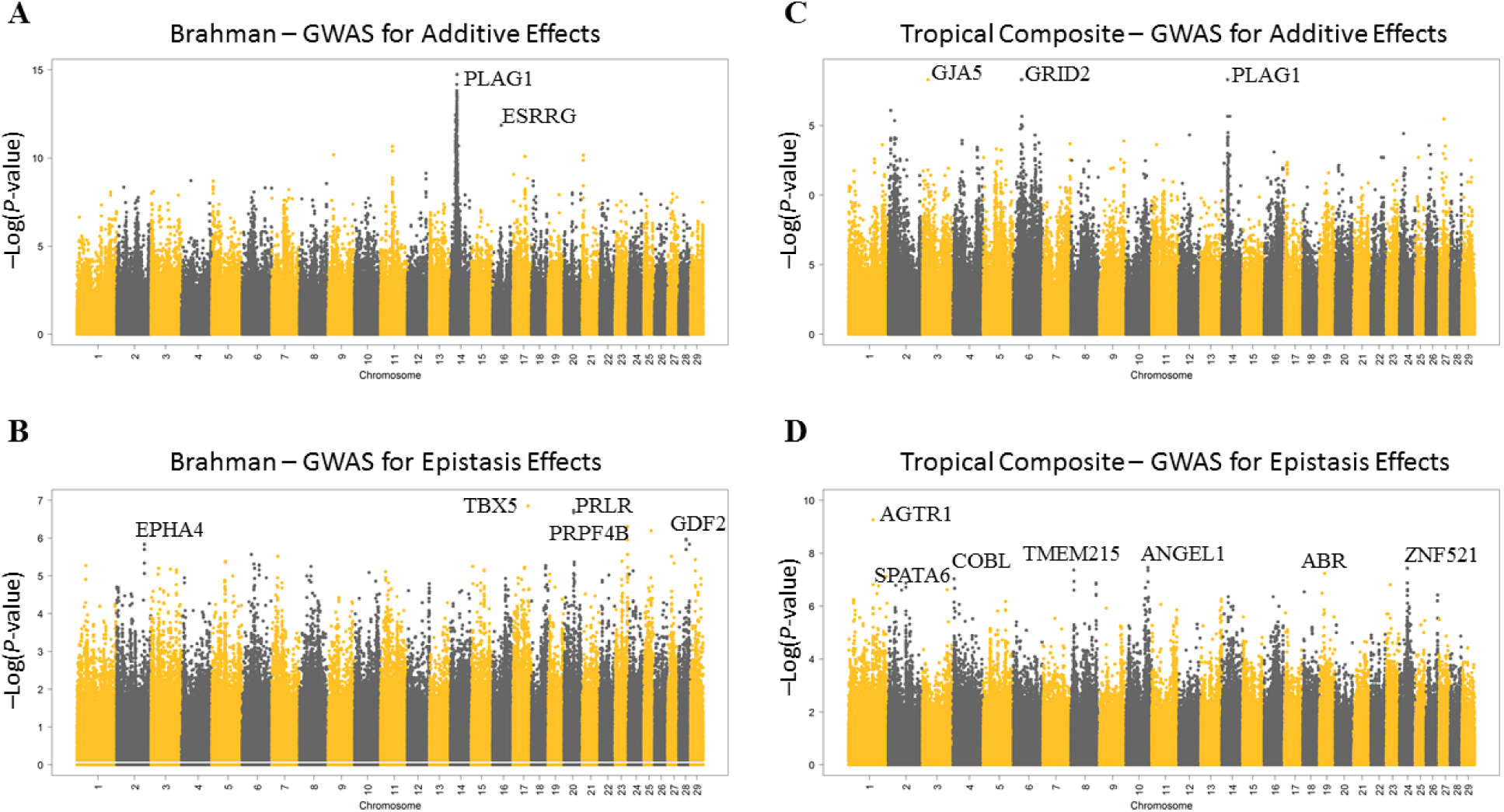
Genome-wide additive and epistasis association: Manhattan plots of the additive (A and C) and epistatic (B and D) association of SNPs across the 29 bovine autosomal chromosomes for the Brahman (A and B) and Tropical Composite (C and D) populations. The most likely candidate genes in the most significantly associated regions are annotated where an obvious candidate could be identified according to the bovine reference genome assembly ARS-UCS1.2. SNPs on odd-numbered chromosomes are in black and those on even-numbered chromosomes are in yellow.

**Figure 2.**
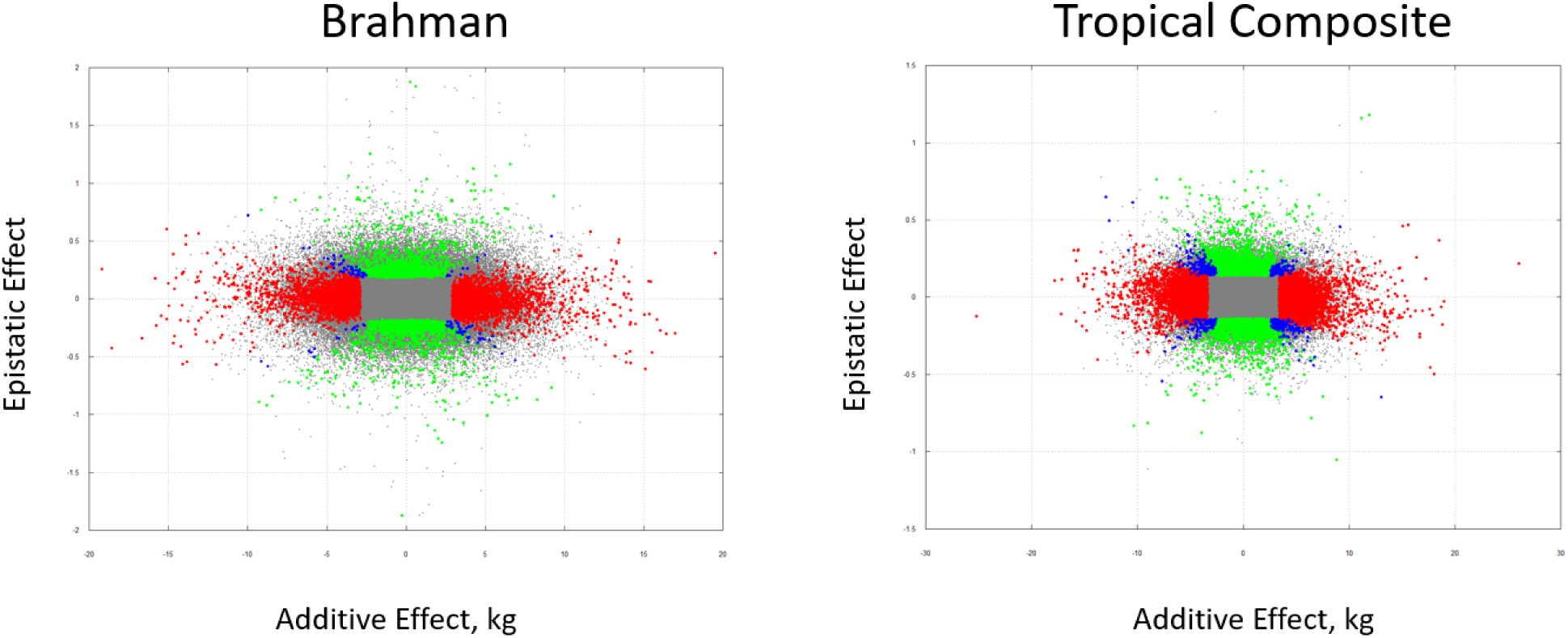
Scatter plot of the relationship between SNP additive (x-axis) and epistatic effects (y-axis) for the Brahman (left panel) and Tropical Composite (right) populations. Red, green and blue indicate significance (P-value < 0.001) for additivity, epistatic and both, respectively.

Figure 1 shows the Manhattan plots for the GWAS for additive and epistatic effects in the BB and TC populations. The most likely candidate gene in the most significantly associated regions are also given in Figure 1. It can already be noted that additive and epistatic gene effects are mutually orthogonal.

In the BB population, the strongest significance for epistatic effect corresponded to SNP BovineHD1700017822 mapped to 60,216,894 bp of BTA17 at 11,627 bp of the coding region of *TBX5* (T-box 5 transcription factor) and with an estimated epistatic effect of −0.533 (-Log10(*P*-value) = 6.849). The corresponding human chromosome segment is involved in ulnar mammary syndrome (Klopocki *et al.* 2006), and a recent large meta-GWAS study reveal TBX5 as a candidate gene for mammary gland morphology in Fleckvieh cattle (Pausch *et al.* 2016).

Following *TBX5*, we found the second strongest signal for epistasis in BB to SNP BovineHD2000011094 (estimated epistatic effect of −0.282 and −Log10(*P*-value) = 6.669) mapped to 38.97 Mb of BTA20 and 2.6 kb downstream of prolactin receptor (*PRLR*). *PRLR* is in a region captured by selection signatures for adaptation in beef cattle (Boitard *et al.* 2016) and mutations on this gene have been found to have a major genetic effect on hair length and coat structure characteristics of cattle (Littlejohn *et al.* 2014; Porto-Neto *et al.* 2018).

The third strongest signal corresponded to SNP BovineHD2300014569 mapped to 50.10 Mb of BTA23 in the coding region of *PRPF4B* (*pre-mRNA processing factor 4B*) with an estimated effect of 0.324 (-Log10(*P*-value) = 6.309). With no reported function in the context of bovine breeding and genetics, *PRPF4B* is an essential kinase induced by estrogen (Lahsaee *et al.* 2016) and its loss promotes sustained growth factor signaling (Corkery *et al.* 2018). Quite strikingly, loci on the coding region of *SPEN* (SNP BovineHD1600014616, epistatic effect = 0.213, −Log10(*P*-value) = 2.137) and *GHR* (SNP BovineHD2000009203, epistatic effect = 0.782, −Log10(*P*-value) = 2.234) were found to be significantly epistatic in our study. *SPEN* is an estrogen receptor cofactor and a key regulator of fat deposition and energy balance (Hazegh *et al.* 2017). Furthermore, a SNP-based co-association gene network by our group previously identified *ESRRG* and *PPARG* as key regulators of age at puberty in Brahman cows (Fortes *et al.* 2013).

In the TC population, we found the strongest signal in SNP BovineHD0100028404 (epistatic effect = 0.294, −Log10(*P*-value) = 9.261) mapped to 98.71 Mb of BTA1 in the coding region of *LOC100139843* (*mCG140927-like*) with limited information known about its function, but quite strikingly, recently reported to be associated with age at puberty in Angus bulls (Fernández *et al.* 2016). We found the second and third strongest signal in the coding region of *ZNF521* (SNP BovineHD2400008618, mapped to BTA24:31,439,030 with and estimated epistatic effect = −0.260, −Log10(*P*-value) = 7.432) and *AGTR1* (SNP BovineHD0100034098 mapped to BTA1: 119,483,491 with and estimated epistatic effect = −0.194, −Log10(*P*-value) = 3.204), respectively. The loci on *ZNF521* has been found to associate with female fertility in Nordic Red cattle, consisting of three different populations from Finland, Sweden and Denmark (Höglund *et al.* 2015). Whereas the role in bovine fertility of *AGTR1* (*angiotensin II receptor type 1*) has long been documented (Portela *et al.* 2008; Marey *et al.* 2016) including its differential expression at the level of the oviduct between *Bos taurus* and *Bos indicus* cattle (Fontes *et al.* 2018).

The relationship between epistatic and additive effect in each population is illustrated in Figure 2. It can be seen that effects are empirically orthogonal as expected. Note that unlike additive effects, epistatic effects have no units as they are defined as a multiplier on additive effects. Figure 3 shows the relationship between the epistatic effects in both populations, BB and TC. Significant simultaneously in both populations were 42 SNPs of which 24 where located within 50 kb of the coding region of known genes and these are listed in Table 2. Porto-Neto et al. (2014) showed that LD dropped below 0.2 at distances of 50 kb.

**Figure 3.**
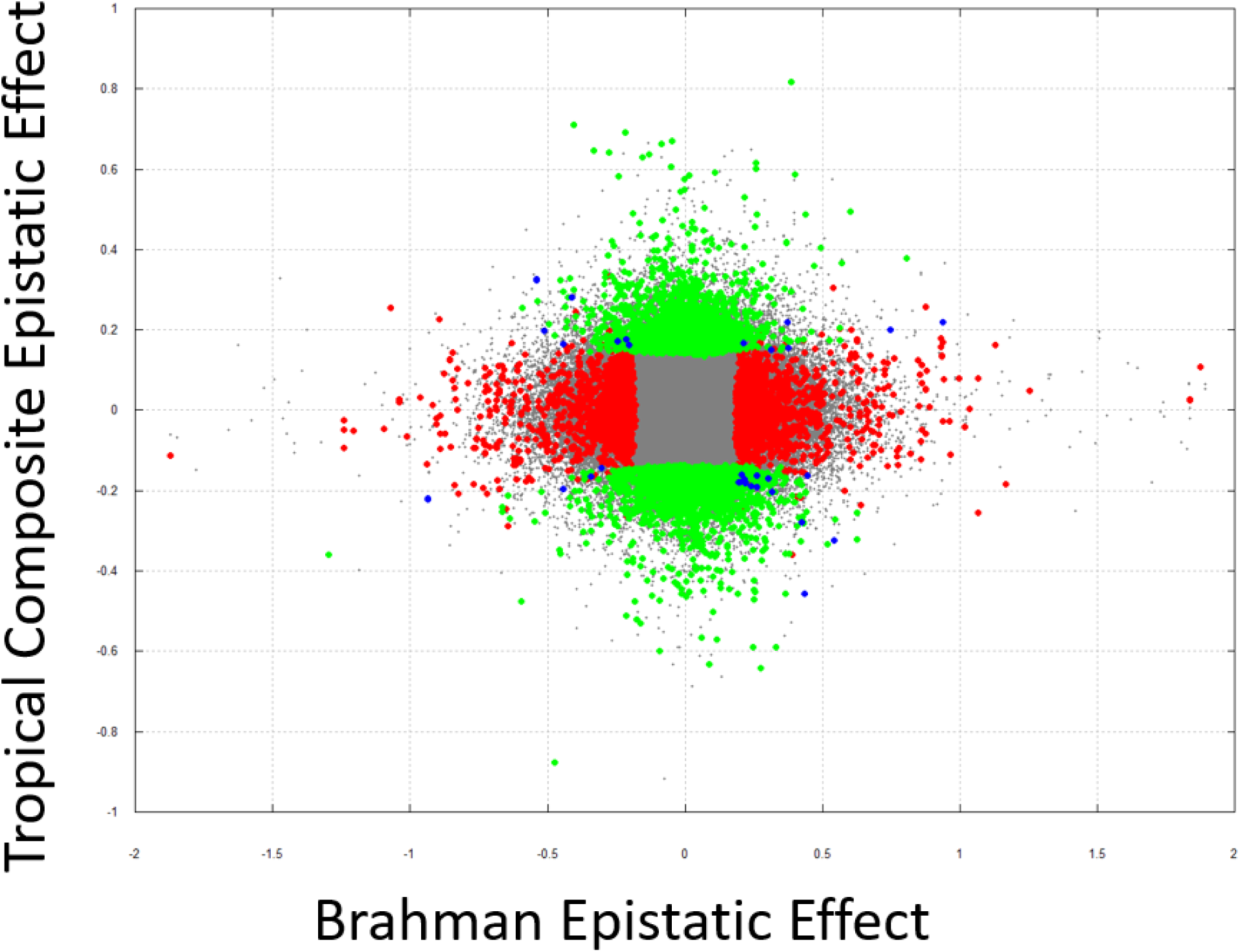
Scatter plot of the relationship between SNP epistatic effect in the Brahman (x-axis) and Tropical Composite (y-axis) populations. Red, green and blue indicate significance (P-value < 0.001) in the Brahman, Tropical Composite, and both populations, respectively.

Among those listed in Table2, prominent genes for their reported role in mammalian fertility including bovine are: *ALDH1A1 (aldehyde dehydrogenase 1 family member A1), PCSK5 (proprotein convertase subtilisin/kexin type 5)*, and *TSHR (thyroid stimulating hormone receptor), and IL21 (Interleukin-21).*

The role of *ALDH1A1* during bovine ovarian development has recently been stablished (Hatzirodos *et al.* 2019; Hummitzsch *et al.* 2019). Antenos et al. (2011) reported the role of PCSK5 in mouse ovarian follicle development. Similarly, *TSHR* is a well-known gene for its function regulating growth, fat metabolism and fertility. Dias et al. (2017) identified a candidate QTL in *TSHR* affecting puberty in five cattle breeds across the taurine and Indicine lineages: Brangus, Brahman, Nellore, Angus and Holstein. Also, one of the most prominent selective sweeps found in all domestic chickens occurred at the locus for *TSHR* (Rubin *et al.* 2010). Finally, the immune system response gene *IL21* has been shown to harbor selection signatures among divergently selected subpopulations of Polish Red cattle (Gurgul *et al.* 2019), and among goats and sheep indigenous to a hot arid environment (Kim *et al.* 2016).

## DISCUSSION

In this study, we present a rigorous formulation of buffering epistasis due to single loci and how it relates to previous works (Jannink 2007; Crawford *et al.* 2017). Then we present an exact test for association analysis of single locus in buffering epistatic interaction based on bi-variate random regression restricted maximum likelihood (REML) analyses.

Methodology for detection of locus by genetic background interactions has been developed at least two times, (sadly) in apparent isolation from each other (Jannink 2007; Crawford *et al.* 2017). Jannink (2007) first and formally derived the linear model involving (additive) genetic background and one locus showing interaction with the background. He showed orthogonality and derived a variance component useful for genome-wide association studies (GWAS), but he did not realize that the model discovered a particular kind of epistasis. Later, the same model was re-derived by Crawford et al. (2017) who, however, did not build an orthogonal model (something that may lead to spurious effect estimates), and could not use maximum likelihood estimates. Neither of the two authors explicitly estimated the effect of the epistatic loci. Here, we complete and interpret their work.

Then we applied our method to two datasets in cattle. Taking together, our findings in these datasets are striking. Some of our significant genes have been reported as having a statistical *additive* effect in other studies, whereas in ours, they have a statistical *epistatic* effect but not an additive one. As shown by theory, a shift in the mean of the genetic values towards one of the extremes changes the statistical effect from epistatic into additive, whereas the functional effect is always epistatic. For instance, a gene with epistatic buffering effect *k*_*i*_ = 0.1 in a population with *ū* = 0 will have a statistically null additive effect, whereas the same gene in a population with *ū* > 0 will have a positive substitution effect, and a negative one in a population with ū < 0. This explains indeed why some effects change sign when observed in different genetic backgrounds (Magwire *et al.* 2010).

A similar argument explains why we find genes found under selection in a multi-breed comparison study. These genes are bound to have little variation in their coding region and/or no additive effect in any given individual breed, because they are fixed or near fixation. However, inside our populations TC and BB, they are identified as relevant because they have opposite effects in the extremes of the polygenic background, and they would get alternative fixation of alleles if the populations were selected towards either extreme. In other words, our epistatic association analysis can be seen as a selection signature analysis *before* selection of the extremes.

It is somehow difficult to grasp the meaning of the estimate of buffering epistatic effects, *e.g.* 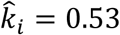 (adimensional) in TBX5 in the BB population. For instance, genetic variation for human height is approximately 39 cm^2^ in European populations (Visscher 2008). Imagine an individual with large total additive genetic value, *i.e.* 4 standard deviations (25 cm taller than the mean, *i.e.* 190 cm for a woman and 203 cm for a man). If such an individual is carrier of the “bufferer” copies *and* the allele has a frequency of 0.8 (we will assume that the bufferer allele is fairly frequent as protective), her/his height changes by *z*_*i*_*k*_*i*_*u* = (2 − 2 × 0.8) × −0.53 × 25, i.e. 5.3 centimeters shorter. Still, the estimate of −0.53 is probably too high – due to the winner’s curse or Beavis effect (Xu 2003).

The possibility of detection and further functional analysis of these epistatic loci implied in buffering mechanisms is of particular relevance in quantitative genetics. As described in the Introduction, these loci must exist (Mackay 2014), be selected (Flatt 2005) and they would explain perplexing phenomena: release or maintenance of additive variance (Visser *et al.* 2003; Flatt 2005), (Gimelfarb 1989), “conversion” of epistatic into additive variance (Mackay 2014) and opposite signs of GWAS associations in different populations (Huang *et al.* 2012) (Magwire *et al.* 2010).

Our model is based on interaction with genetic background, and there is increasing evidence of dependency of gene substitution on genetic background (Hansen 2013). This implies, potentially, large changes in gene effects and selection dynamics even if genetic variance is nearly fully additive at each step (Hansen 2013). Indeed, it has been argued that epistasis provides the basis of rapid adaptation to new environments (Wright 1931; Mackay 2014).

For evolutionary research and plant and animal breeding, there is growing interest in understanding the extent and mechanisms of epistasis in biology because of the intriguing prospect of it being an untapped future source of additive variation that may be exploited by nature and by breeding programs to evolve phenotypes as well as influencing genetic heterogeneity. Indeed, the role of epistasis in determining which mutations ultimately succeed or fail in a population under selection remains a central challenge in biology. Buskirk et al. (2017) demonstrated the power of experimental evolution to identify epistatic interactions. In multi-generation selection programs continued response has been seen, classically for over 100 generations in the Illinois maize kernel content lines, and there have been large and still continuing genetic improvements in livestock populations, notably in broiler chickens (Hill 2016). Whilst the obvious source of continued response is *de novo* mutation, some of the additive variation being utilized may have derived from existing mutations whose behavior changes from epistatic to additive in response to changes in the remainder of the genome, consistent with the argument of Carlborg et al. (2006). Paixão and Barton (2016) argued that ‘epistasis sustains additive genetic variance for longer: Alleles that were initially deleterious or near-neutral may acquire favorable effects as the genetic background changes, “converting” epistatic variance into additive, and so prolonging the response to selection’. Similarly, Hill (2017) concluded that ‘It seems better to concentrate on utilizing additive variance, and hope for a bonus from converting epistatic variance’.

To conclude, our method and our results show promising new perspectives for research in evolutionary and quantitative genetics based on the study of loci that buffer effect of other loci.

## ACKNOWLEDGMENTS

Excellent advice and review was provided by Sonja Dominik and Andrew George. Work supported by INRA SELGEN metaprogram (projects EpiSel and OptiMaGics) and INRA-CSIRO Linkage Action and SMARTER. Project partly supported by Toulouse Midi-Pyrénées bioinformatics platform.

## AUTHORS’ CONTRIBUTIONS

AR, JH, KM, ZV and AL derived the equations, wrote the programs to do the analyses, performed analyses, and assisted drafting the manuscript. L P-N and M N-S provided sequence-level data, and designed and performed the functional analyses. FR, YL and NH provided valuable insights throughout the analysis and writing process.

### APPENDIX

#### Technical details for the REML analyses

Because the covariance matrix in the alternative model is not full rank,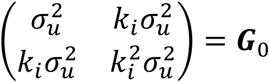, the reduced rank method of (Meyer and Kirkpatrick 2005) was used (explicitly stating rank equal to 1), and the iterative algorithm was a mixture of PX and AI (we refer the reader to the Wombat manual for details). Unlike other software programs, Wombat does not accept centered genotypes as covariates for the random regression model – only integers *m* = {1,2,3} for each genotype that are transformed into reals (centered and scaled) internally. Thus, although the likelihood is in the correct scale, the estimated 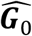 on output is not. In a model with random regressions, ***y*** = … + ***Pu*** + … the covariance across two points is a function of ***PKP***′ where ***K*** = *Var*(***u***) is a matrix of covariances and ***P*** are covariates. Accordingly, to put back 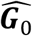 in the right scale we need to construct, for each locus *i*, 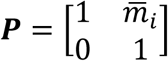 where 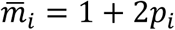 is the mean of the regressors *m* = {1,2,3}. Back transformation to the regular scale is carried out using 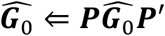 and from here the correct value 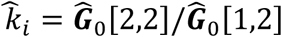.

